# The exchange rates of amide and arginine guanidinium CEST in the mouse brain

**DOI:** 10.1101/2022.02.14.480399

**Authors:** Kexin Wang, Ran Sui, Lin Chen, Yuguo Li, Jiadi Xu

## Abstract

**Purpose:** To develop a pipeline for measuring the exchange rates and concentrations of in vivo excgangeable protons, and to demonstrate this for the amide and arginine (Arg) guanidinium (Guan) protons in mobile proteins in the mouse brain.

**Methods:** An ultra-short echo (UTE) CEST sequence with a continuous wave presaturation (preRadCEST) was applied to acquire Z-spectra with robustness to motion and physiological fluctuations. AmideCEST and Arginine guanCEST (ArgCEST) were extracted and their proton concentrations and exchange rates obtained using a two-step multi-B_1_ Bloch fitting approach that included the semisolid macromolecular background. To minimize contamination from the amine protons from creatine and phosphocreatine, ArgCEST measurements were performed on the Guanidinoacetate N-methyltransferase deficiency (GAMT^-/-^) mouse characterized by low creatine and phosphocreatine concentrations in the brain.

**Results:** For the amideCEST proton pool, the exchange rate and concentrations were found to be 59.6 ± 9.0 s^-1^ and 41.7 ± 7.0 mM, respectively, with the maximum signal observed at B_1_ = 0.8 μT. For the ArgCEST proton, the guanidinium exchange these were 70.1 ± 5.5 s^-1^ and 10.1 ± 1.3 mM, respectively, with the maximum effect observed at B_1_ = 0.9 μT. The current study suggests that the inverse pH dependence in GuanCEST of brain is led by the CrCEST component, not ArgCEST.

**Conclusion:** The current pipeline is expected to have general use for *in vivo* CEST quantitation and optimization of visible CEST resonances.

## Introduction

Chemical exchange saturation transfer (CEST) MRI is a molecular imaging method that can significantly enhance the sensitivity of detecting low concentrations of proteins and metabolites through the water signal (1-3). Among many endogenous CEST contrasts, amide protons (amide proton transfer, APT, or amideCEST) in mobile proteins have been intensively studied. AmideCEST signal has a well-defined lineshape, which stands out in the crowded *in vivo* Z-spectrum (4,5). It is noteworthy that amideCEST should be distinguished from APT-weighted MRI, which also focuses on the peak at 3.5 ppm but is obtained by asymmetry analysis of the Z-spectrum and thus includes multiple other effects including those from relayed NOEs (rNOEs) and asymmetry in the semisolid magnetization transfer contrast (MTC) (6). Guanidinium (Guan) CEST was reported in creatine (Cr) phantoms (7-9), muscle (10-13), and brain (14), the Z-spectrum of which at 2 ppm also includes the phosphocreatine (PCr) and the guanidinium protons of arginine groups in endogenous mobile proteins (5,15-18). Since for proteins the only guanidinium group exists in Arginine (Arg) groups, we name the GuanCEST signal from proteins as ArgCEST, while Cr as CrCEST. AmideCEST and ArgCEST are among a few CEST contrasts that can be extracted with high confidence from the crowded *in vivo* Z-spectrum because of their distinguishable peaks at multiple MRI field strengths. The optimum acquisition and quantification of amideCEST and ArgCEST contrasts usually requires knowledge of their exchange rates. Particularly, the exchange rate of the ArgCEST proton pool is critical in the understanding of the pH and B_1_ dependence of signal at 2 ppm (15,19-21). For most metabolite CEST studies, the exchange rates can be measured in phantom solution under physiological conditions, such as glutamate (22), D-glucose (23,24), Cr (9), and PCr (9,25,26). However, this method is not possible for both amideCEST and ArgCEST since these resonances reflect the composite effect of multiple unidentified proteins. An initial attempt to determine the amide proton exchange rate was carried out in the rat brain with a water-exchange filter (WEX) NMR sequence, giving a value of 28.6 s^-1^ (27). Recent studies, including frequency labeled exchange transfer MRI (28) and fingerprinting CEST method, suggested a much higher exchange rate (around 200-400 s^-1^) (29), while another group reported as 34.6-47.9 s^-1^ with fingerprinting (30,31). Hence, it is worth revisiting the amideCEST exchange rate with recent technology and also fill in the long-standing blank of the ArgCEST exchange rate.

The direct measurement of exchange rates of amideCEST and ArgCEST from the brain Z-spectrum is hampered by three factors. Firstly, the acquisition of Z-spectrum needs correction for the noise and artifacts introduced by physiological motions (32,33) and/or MRI instability. A large variation is observed in the Z-spectrum when the saturation field strength in Hertz is comparable to the B_0_ fluctuation in the mouse brain caused by the respiration and brain movements (34). This can be partially improved using the motion-insensitive CEST MRI methods developed recently, such as the self-gated ultra-short echo time (UTE) CEST technique (35-37), or using navigators to correct for frequency fluctuations and motion. (38-40) Secondly, the extraction of both amideCEST and ArgCEST signals from *in vivo* Z-spectrum is challenging. Multi-pool Lorentzian line-shape fitting does not function well for the amide and guanidinium resonances (41-45), due to their overlap with amineCEST, PCr amide, and amide/aromatic rNOE signals, especially for guanidinium. Fortunately, recent pH-dependent studies on animal stroke models at both high (4,15,21) and low MRI fields (18) have suggested that both amideCEST and ArgCEST show discernible peaks with smooth backgrounds in the brain Z-spectrum. These peaks can be well extracted and quantified using a recently proposed polynomial and Lorentzian line-shape fitting (PLOF) approach (17,20,25,34). Another challenge for the ArgCEST measurement is the contamination from Cr guanidinium and PCr amide peaks around 2 ppm (16,17). This problem can be solved by performing the ArgCEST assessment using a Guanidinoacetate N-methyltransferase deficiency (GAMT^-/-^) mouse model that has low Cr and PCr concentrations in the brain (46,47). Thirdly, the quantitation of the CEST exchange rate in the brain with the QUESP (the quantification of exchange rate using varying saturation power) (48,49) and Omega plot (50-52) methods needs to account for the strong MTC and direct saturation backgrounds in tissues. To overcome all of the challenges mentioned above, we developed a two-step multi-B_1_ Bloch-McConnell (BM) fitting approach for both signal extraction (background removal) and quantification. The idea is similar to that behind PLOF CEST, but strictly follows the BM equations for fitting the MTC background and the CEST peaks. The approach was first validated in a creatine (Cr) phantom mixed with cross-linked bovine serum albumin (BSA) and then applied *in vivo* for the amideCEST with wild type (WT) mouse and ArgCEST using GAMT^-/-^ mouse brains.

## Methods

### Presaturation Radial CEST sequence

The CEST pulse sequence with a radial readout (35-37,53) was composed of a train of interleaved Gaussian saturation pulses and 2D-UTE modules. As demonstrated in both brain (35) and liver CEST studies (36), a radial acquisition scheme is inherently less sensitive to motion artifacts. However, the steady-state UTE-CEST is difficult for CEST quantification as it is challenging to model the pulsed CEST signal. Therefore, we modified the sequence to perform continuous wave (CW) CEST labeling at the beginning followed by a train of 2D UTE readout modules as shown in Fig. 1, which is dubbed presaturation radial CEST (preRadCEST).

**Figure 1.**
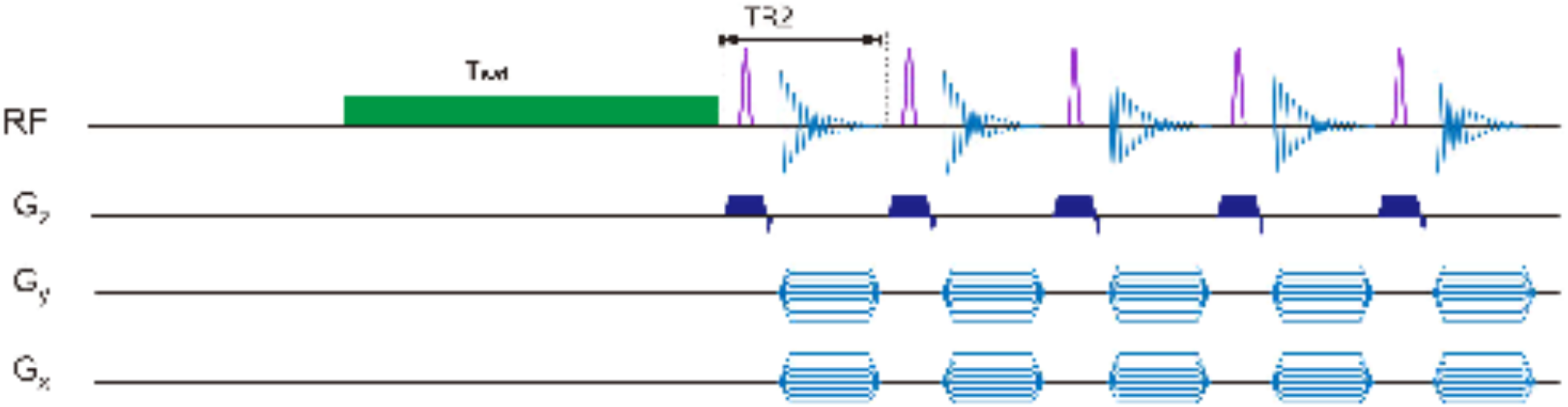
(a) Timing diagram of the preRadCEST sequence. T_sat_ is the length of the CW saturation period, while TR2 is the time duration between successive readout pules.

### CEST extraction and quantification

Three pools, i.e., water, magnetization transfer contrast (MTC) and CEST, were assumed for the BM equations. Here, MTC is a combined pool from both macromolecules and other CEST components. For the macromolecular MTC proton pool, the transverse relaxation time can be neglected due to its extremely short T_2_ (∼10 μs) (54). Then, the MTC components are determined by three parameters (supplemental material Eq. S4): the MTC exchange rate *k*_MTC,_ the back exchange rate from MTC to water *k*_*MTC*_·*f*_*MTC*_, and the *R*_1*MTC*_+*R*_*MTC*_. *R*_1*MTC*_ is typically much lower than *R*_*MTC*_ and has minimum impact on the fitting, and was set to 1 s^-1^ here. *T*_2MTC_ usually determines the MTC lineshape in the conventional Super-Lorentzian or Gaussian assumptions for the biological tissues (55,56). In current study, a linear lineshape was used for the mixed MTC and

CEST background as we fit only a small range of the Z-spectrum (< 4 ppm). Then, *T*_2MTC_ has no impact on the fitting results and was set to 10 μs.The detailed BM fitting is provided in the supporting information and the Matlab code for the BM fitting will be made available at https://github.com/jiadixu/Two-step-Bloch-Fitting after the paper is accepted. The Z-spectra acquired with different B_1_ values are fitted simultaneously, i.e., group BM fitting. Two steps of the fitting are performed as below:

1. The MTC and water background spectrum were fitted by excluding the data points for amideCEST (or ArgCEST) with varying parameters, *T*_*2app*_, 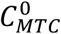 and 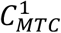, for each Z-spectrum. Here, *T*_*2app*,_ is a parameter including water transverse relaxation and the broadening caused by the fast-exchanging protons. The offset was fixed at 3.5 ppm (amideCEST) and 2.0 ppm (ArgCEST), while T_1_ of the amide and ArgCEST pool was set to 1 s. The *T*_*1w*_ values measured in both phantom and mouse brain were fixed for the group fitting. The fitting was achieved by minimizing the root mean square (RMS) of the difference between our fitting and experimental data. For amideCEST and ArgCEST, background fitting ranges of 2.0–5.5 ppm and 1.0-3.5 ppm were used, excluding the relevant signal ranges of [3.2, 4.0] ppm and [1.6, 2.6] ppm, respectively.
2. The extraction and fitting of amideCEST and ArgCEST signals were achieved by fixing the background signal obtained in step 1, i.e., the *T*_1w_, *T*_*2app*_, 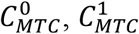 values. The *T*_2amide_, *k*_amide_ and *f*_amide_ (or *T*_2Guan,_ *k*_Guan_ and *f*_Guan_) values were varied in the amideCEST (or ArgCEST) fitting to minimize the RMS of the difference between our fitting and experimental data. Either amideCEST or ArgCEST signal can be obtained by subtracting the background signal from the CEST signals at 3.5 and 2.0 ppm respectively.

### Phantom Validation

The two-step BM fitting method was validated on a Cr (50 mM) solution mixed with 15% cross-linked BSA (CrCrossBSA) in phosphate buffered saline (PBS), titrated to pH 7.7. BSA crosslinking was achieved using a 25 μL glutaraldehyde solution. The exchange rate of the CrCrossBSA phantoms was measured on a Bruker 17.6T NMR spectrometer using the selective inversion recovery method detailed in the supplemental material (13).

### MRI Experiments

Animal procedures were approved by the Johns Hopkins University Animal Care and Use Committee. Six C57BL/6 mice (age: 3 months) and four GAMT^-/-^ mice (age: 3 months) were used. All MRI experiments were performed on a horizontal bore 11.7 T Bruker Biospec system (Bruker, Ettlingen, Germany). A 72 mm quadrature volume resonator was used as a transmitter, and a 2×2 phased array was used as a receiver for both phantoms and animals. Animals were anesthetized using 2% isoflurane in medical air, followed by 1% to 1.5% isoflurane for maintenance. The B_0_ field over the image slice was adjusted to second order for animal using field-mapping and shimmed on the water linewidth for phantoms. T_1_-weighted maps were acquired using the same geometry and spatial resolution as CEST MRI, employing a RAREVTR sequence (RARE with variable TR = 0.5, 1, 1.5, 2, 3.5, 5, 8 s).

The pre-saturation time was set to 2 s for all CEST experiments. Seven B_1_ values were collected for the CrCrossBSA phantom (0.4, 0.6, 0.8, 1.0, 1.2, 1.4 and 1.6 μT), while eight B_1_ values were used for the mouse brain (0.4, 0.5, 0.6, 0.8, 1.0, 1.2, 1.4 and 1.6 μT). The frequency was swept from 2.0 to 5.5 pm with an increment of 0.1 ppm for the amideCEST. This same frequency step size was applied for the ArgCEST from 1.0 ppm to 3.5 ppm, except for a smaller increment of 0.05 ppm between 1.4 ppm and 3.0 ppm. For phantom studies, the frequency was swept from 0.5 to 3.5 ppm with 0.1 ppm increment. The excitation pulse for the 2D-UTE radial readout was 0.4 ms with a flip angle of 10 degrees. After each saturation pulse, one radial spoke was collected with TR2 = 6 ms. In the current study, single-slice CEST images were collected with a total acquisition time of 900 ms. The nominal Cartesian matrix size was set to 96×96 with the slice thickness = 2 mm. Before performing experiments on the GAMT^-/-^ mice, *in vivo* MRS was performed on the mouse brain to confirm the low concentration of Cr/PCr (Supporting Information Figure S1).

## Results

### Phantom validation experiments

One challenging part in the BM fitting is the great number of parameters that participate, while they are interacting with each other. We used a CrCrossBSA phantom as ground truth to validate the fitting method. The typical MRS spectrum of CrCrossBSA is plotted in the inset of Fig. 2a. The Guan exchange rate at room temperature and pH 7.7 was determined to be 106 ± 8 s^-1^ by fitting the selective inversion curves of the Guan peak (Fig. 2a).

**Figure 2.**
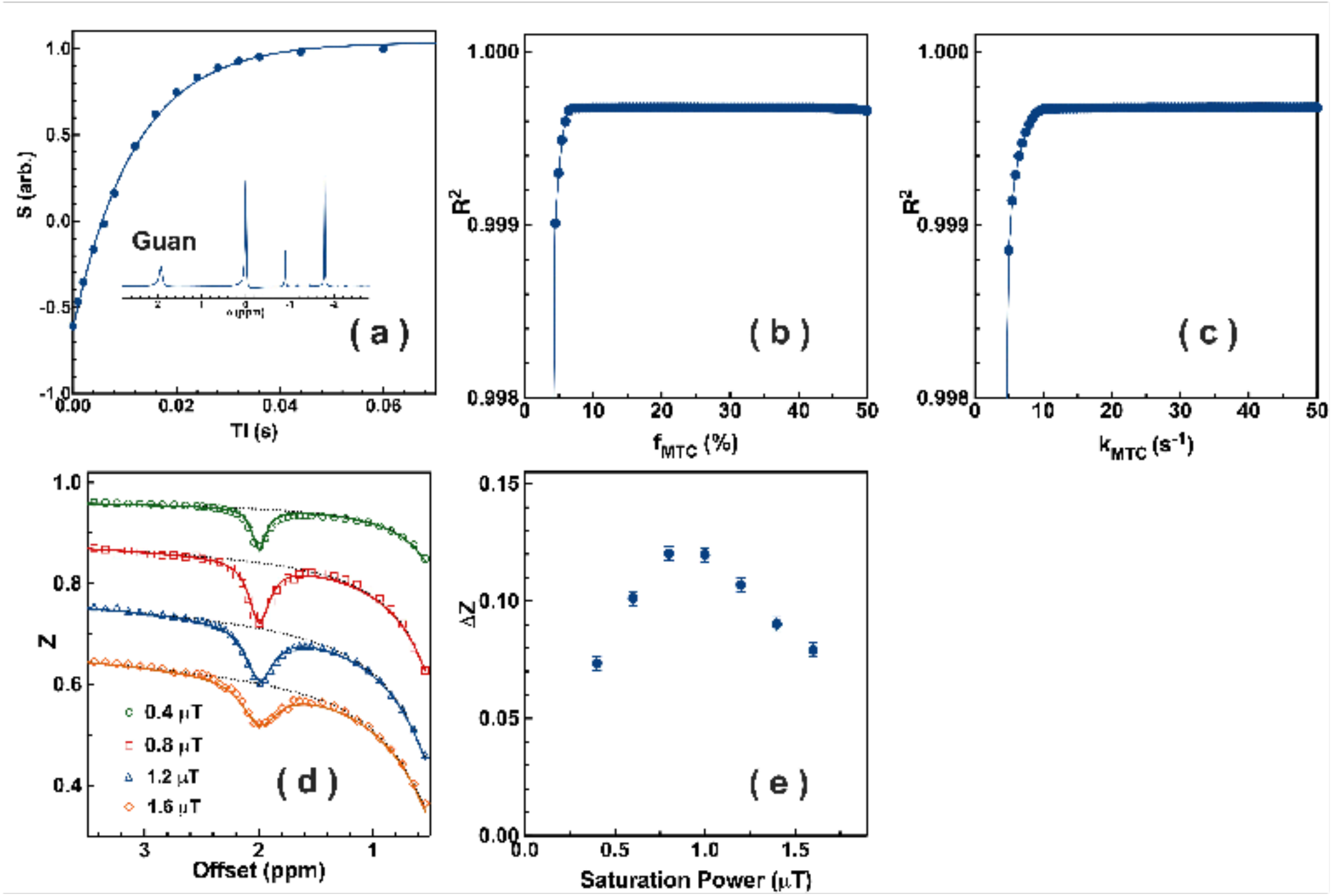
(a) The inversion recovery curve of peaks at 1.92 ppm (Guan) of Cr in 15% cross-linked BSA at room temperature and pH 7.7. The curve was fitted using a single exponential curve (Supporting Information Eq. S1). Inset: typical 1D NMR spectrum of CrCrossBSA phantom. (b-d) The impact of different (b) *f*_MTC_ and (c) *k*_MTC_ on the MTC background fitting. The goodness of the fitting results was evaluated by the R^2^ coefficient of determination. (b) *k*_MTC_=20 s^-1^ set for the simulation (c) In the *k*_MTC_ simulation, *f*_MTC_=10%. (d) Typical Z-spectra of the GuanCEST region recorded with a B_1_ of 0.4, 0.8, 1.2 and 1.6 μT and an illustration of the two-step multi-B_1_ BM fitting scheme for extracting and quantifying the GuanCEST signal (R^2^=0.995). All CrCrossBSA experiments were performed at room temperature. The black dashed lines are the background Z-spectrum, while the solid lines are fitted CrCEST curves with the fixed water and MTC background. In the whole fitting, the measured water *T*_*1w*_ = 1.8 s was used, while *T*_1MTC_ = 1 s and *T*_2MTC_ = 10 μs for the MTC pool were fixed. (f) The B_1_-dependent CrCEST values for the CrCrossBSA phantom extracted with the two-step multi-B_1_ BM fitting.

CEST experiments were performed on the same phantom at room temperature with the preRadCEST sequence. To examine the proper selection of *k*_MTC_, *f*_MTC_ and *T*_2MTC_ values on the MTC background fitting. The MTC background fitting in the step one was evaluated with different *k*_MTC_, *T*_2MTC_ and *f*_MTC_ values while *T*_*2app*_, 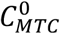 and 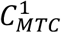 varied for each B_1_ Z-spectrum (Figs. 2b-d). The goodness-of-fit was evaluated by the *R*^*2*^ coefficient. The result suggested that the fitting is insensitive to the *f*_MTC_ and *k*_MTC_ when *f*_MTC_ > 8% and *k*_MTC_ > 11%. Hence, typical in vivo *k*_MTC_ = 20 s^-1^, *f*_MTC_ = 10% were fixed in our study, which are representative *in vivo* MTC parameters determined by on- and off-resonance variable delay multi-pulse (VDMP) methods (57,58) as well as previous MTC studies (59-65)

The typical Z-spectrum as well as the BM fitting curves are plotted in Fig. 2e. The fitting curves matched the observed Z-spectrum for all B_1_ values (R^2^ > 0.995). From the two-step BM fitting, CrCEST exchange rate was determined to be 104.7±6.0 s^-1^, concentration = 46.8±5 mM and *T*_2Guan_ = 18.3±2 ms. The exchange rate and concentration matched well with the measured rate from the inversion experiment (106 ± 8 s^-1^) and the prepared concentration (50 mM). The fitting results for all parameters are listed in the supporting information Table S1. The CrCEST signal as a function of B_1_ plotted in Fig. 2f shows typical buildup and decay pattern with the peak at 0.8 μT.

### AmideCEST in the WT mice

In Fig. 3a, the averaged (n=4) amideCEST Z-spectrum of the WT mouse whole brain is shown for four typical B_1_ values. The curve fitted using the two-step BM fitting method demonstrates the extraction of the amideCEST and the high quality of fitting. The amideCEST signals extracted by our fitting are plotted as a function of B_1_ in Fig. 3b. An exchange rate of 59.6 ± 9 s^-1^, a concentration of 41.7 ± 7 mM and *T*_2s_ = 5.0 ± 0.9 ms were extracted from the fitting.

**Figure 3.**
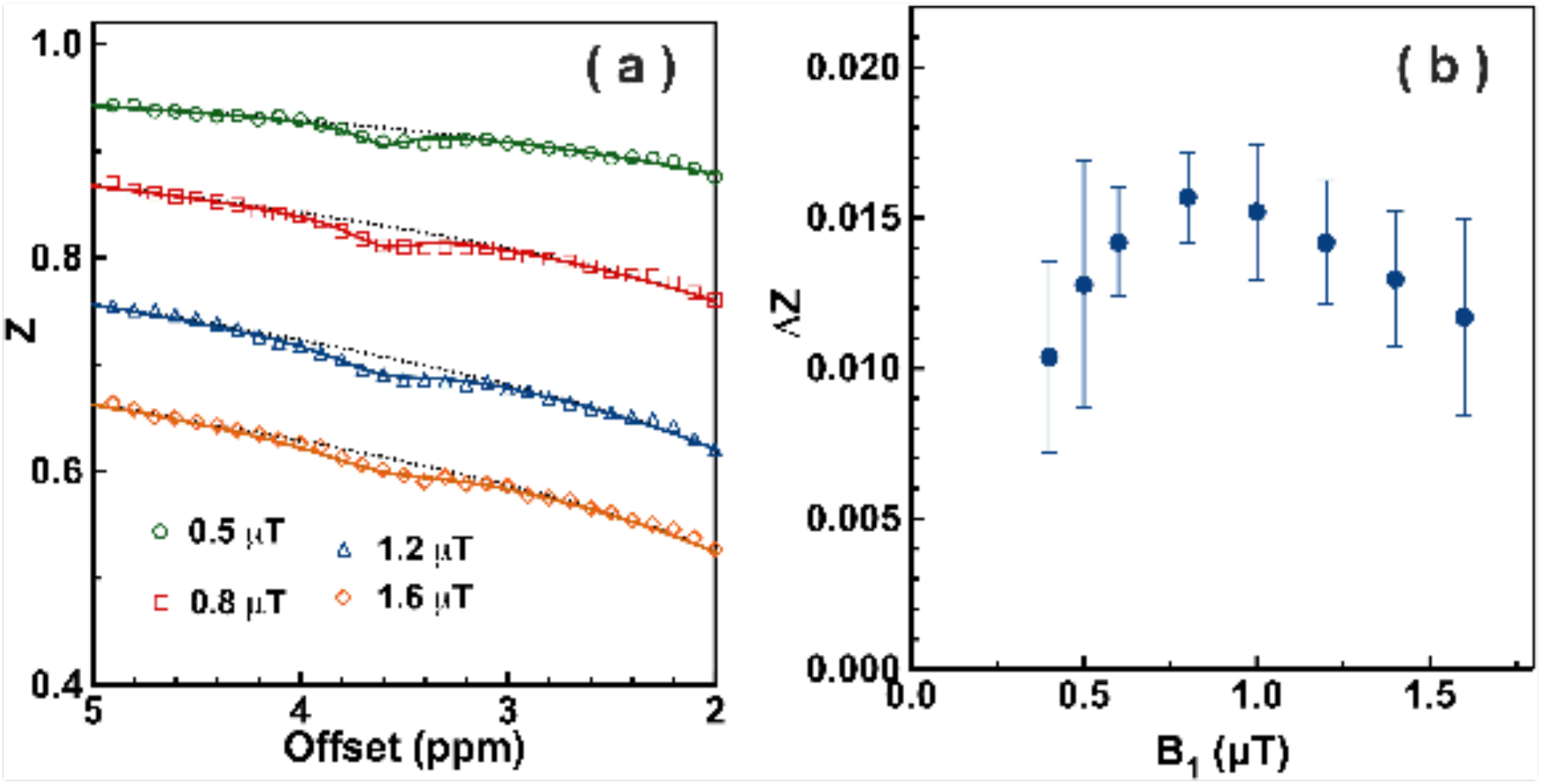
(a) Averaged Z-spectrum of the amideCEST (n=4) for the WT mice recorded using preRadCEST with a fixed saturation length of 5 s and varying B_1_ =0.5, 0.8, 1.2 and 1.6 μT together with the two-step B1 group fitting curves. Dashed lines are the MTC background fitting curves, while solid lines are the difference Z-spectrum fitting. (b) The extracted amideCEST signal (ΔZ) of the WT whole brain as a function of B_1_ extracted with the two-step BM fitting.

### ArgCEST measurement in the GAMT^-/-^ mice

The averaged (n=4) Z-spectrum of the GAMT^-/-^ mouse whole brain of is shown in Fig. 4a for the ArgCEST region. The fitted curves using the two-step group multi-B_1_ BM fitting method are also displayed to demonstrate the MTC background fitting results and the extraction of the ArgCEST effect. An exchange rate of 70.1 ± 5.5 s^-1^, a concentration of 10.1 ± 1.3 mM and *T*_2s_ = 3.9 ± 0.5 ms were extracted from the fitting for the *in vivo* ArgCEST. The extracted MTC background parameters *T*_*2app*_,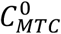, and 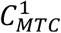 as a function of 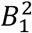 are plotted in Fig. 4b-d. They can be well fitted with linear functions (R^2^>0.83) and are given by

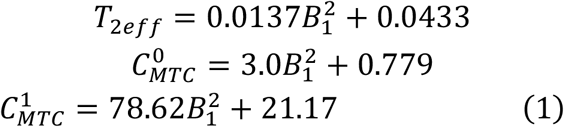

The above equations were used for the following B_1_ dependent CrCEST and ArgCEST simulation. In Fig. 4e, the averaged ArgCEST signal extracted using two-step BM fitting with respect to the saturation power is plotted with the BM simulation curves at fixed MTC background parameters. The optimum saturation power for ArgCEST is 0.8 μT according to Fig. 4e. The CrCEST signals with different exchange rates, i.e., 150, 400 and 1000 s^-1^ were also simulated by fixing the concentration at 5 mM. With exchange rates between 150-400 s^-1^, the curves show clear buildup and decay patterns with peaks at 1.0-1.4 μT, while the signal is buildup and the signal hardly decays at a high exchange rate of 1000 s^-1^. To explain the pH dependent CEST contrasts, the ArgCEST signal as a function of exchange rate was simulated with BM equations for four typical B_1_ values (0.5, 1, 2 and 3 μT) displayed in Fig. 4f by fixing the MTC background.

**Figure 4.**
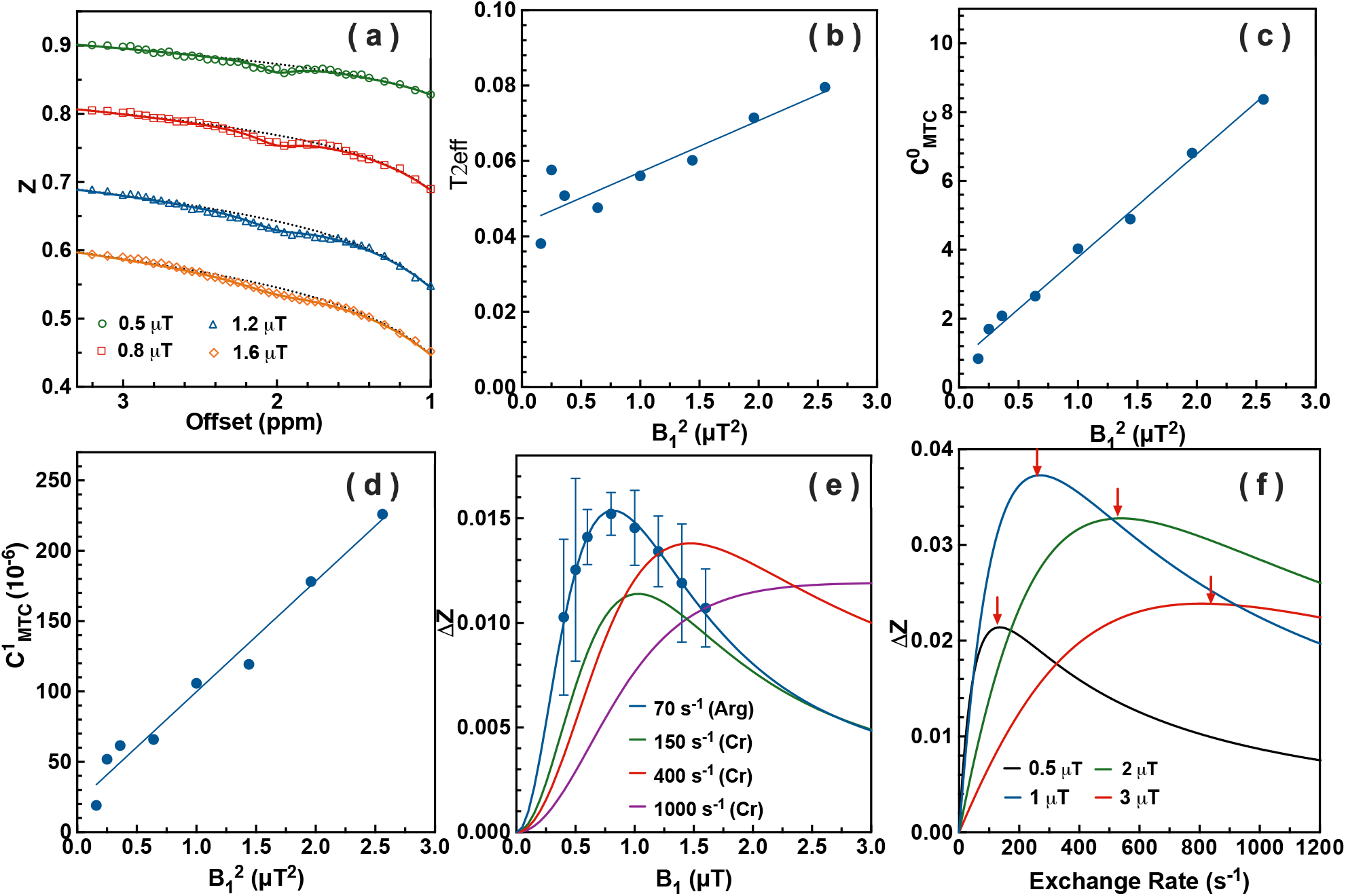
(a) Averaged Z-spectrum of the ArgCEST (n=4) for the GAMT^-/-^ mice recorded with four typical B_1_ values together with the two-step BM fitting curves. Dashed lines are the MTC background fitting curves, while solid-lines are the difference Z-spectrum fitting. (b-c) The extracted MTC background parameters (b) *T*_*2app*,_ (c) 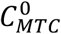, and (d) 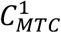 as a function of 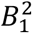. Solid lines are the linear fitting curves. (e) The averaged saturation power-dependent ArgCEST values at 2 ppm for the whole brain of GAMT^-/-^ mouse (n=4). The solid lines are the simulated curves for both Arg and Cr GuanCEST with different exchange rates and concentrations, i.e., 70.0 s^-1^ (10.0 mM) for Arg, 150 s^-1^ (5 mM), 400 s^-1^ (5 mM) and 1000 s^-1^ (5 mM) for Cr. In the fitting, *T*_2s_=3.9 ms, *T*_1w_=1.8 s, *T*_1MTC_=1 s, *T*_2MTC_=10 μs, *f*_MTC_=10%, saturation time 5 s was fixed. (f) The simulated exchange rate dependent GuanCEST signal for four typical saturation powers, i.e., 0.5 μT, 1 μT, 2 μT and 3 μT. The Rabi frequencies are labeled with red arrows, i.e. 134 s^-1^ (0.5 μT), 268 s^-1^ (1 μT), 538 s^-1^ (2 μT), 806 s^-1^ (3 μT). In the fitting, *f*_guan_=11.9 mM, saturation time 5 s and *T*_2guan_=15 ms were used.

## Discussion

In this paper, we used recently developed CEST acquisition, extraction, and quantification methods to determine the exchange rates and concentrations of both amide protons and guanidinium protons in arginine residues of mobile proteins in the mouse brain. Using a t_sat_ of 2 s, the maximum amideCEST signal was observed with a B_1_ of 0.8 μT, comparable to the previous observed value (1 μT) with a three-point method (4). The measured amide exchange rate of 59.6 s^-1^ was slightly higher than that measured previously by the WEX method (28.6 s^-1^) (27), but is consistent to values measured with some of the MR fingerprinting methods (34.6-47.9 s^-1^) (30,31). The consistent results from all three methods further confirm these methods are measuring the same protons, while other methods may have used acquisition parameters that are more sensitive to mixed effects of other proton pools possibly increasing the determined exchange rates.

Previous power dependent studies on both brain and muscle suggested an optimum B_1_ of 1.0-1.4 μT for *in vivo* CrCEST (17,25,26), which indicates the *in vivo* CrCEST exchange rate is far lower than the exchange rates in solution (950 s^-1^) by comparing with simulations in Fig. 4e. The CrCEST exchange rate in tissue is estimated between 150-400 s^-1^ from the optimum B_1_ and rapid signal decay with B_1_>1.4 μT in the in vivo studies (17,25,26). The Cr exchange rate in solution was estimated to around 1000 s^-1^ at room temperature and pH 7.7 (66). The CrCEST exchange rate in the cross-linked BSA is significantly lower than that of the Cr solution (106 s^-1^ vs 1000 s^-1^). The exact reason for the much lower Cr exchange rates in both tissues and cross-linked BSA is not clear. The exchange rate of ArgCEST (70.1 s^-1^) found here is far smaller than that of CrCEST in the physiological pH range, which should allow the selective enhancement of the CrCEST signal in the WT mouse brain by applying high B_1_ (Fig. 4e).

The measured guanidinium proton exchange rate determined from ArgCEST also reveals the reason behind the opposite pH-dependence of ArgCEST signal under different conditions of B_1_ on mouse brain. It is known that the CEST signal increases with decreased exchange rates when the exchange rates are higher than the Rabi frequency of the saturation pulses used (see Fig. 4f) (67,68). The guanidinium exchange rate in Cr (150-400 s^-1^) is above the Rabi frequency (134 s^-1^) corresponding to B_1_ = 0.5 μT and should show inverse dependence on the exchange rate, i.e., pH, as demonstrated in the Fig. 4f. On the hand, the ArgCEST (70.1 s^-1^) is still lower than the Rabi frequency (134 s^-1^) and should be proportional to pH for this B_1_ (Fig. 4f). Therefore, the current study suggests that the inverse pH dependence in GuanCEST is led by the CrCEST component as the GuanCEST in WT mouse brain is a combined signal of CrCEST and ArgCEST (15). Although the protamine ArgCEST was also found to be inversely dependent on pH with low B_1_ (15), this observation cannot be used to validate the *in vivo* pH dependence due to its extremely high exchange rates as shown in the Supporting Information Figure S2. The exchange rate of Arg guanidinium protons in protamine solution is 743 ± 20 s^-1^ at pH 7.0. With high saturation RF strengths such as 2 μT, both Arg and Cr guanidinium exchange rates are lower than the Rabi frequency (532 s^-1^) and hence proportional to pH. This explains the strong pH dependence found in mouse brain with B_1_ = 2.0 μT (20,21). The slow exchange rate of ArgCEST is also consistent with the observation in eggwhite at 3T, in which a clear ArgCEST peak is visible (5).

## Conclusion

We used preRadCEST acquisition, two-step group BM extraction and quantification for determination of the exchange rates and concentrations of amideCEST and ArgCEST in the mouse brain. This approach can be generalized for other CEST quantitation and optimizations as well as *in vivo* CEST ratiometric approach for pH imaging.

## Acknowledgment

The authors thank Drs. Dirk Isenbrandt, Robert Weiss, and Michelle Leppo for creating and providing the GAMT^-/-^ mice.

## Data Availability Statement

The code that supports the findings of this study will be made available at https://github.com/jiadixu/Two-step-Bloch-Fitting.

## Supporting Information

### Bloch-McConnell (BM) Simulations

The observed MTC/CEST signal under influence of a continuous RF pulse can be calculated using the coupled Bloch equations for a three-pool model: water (*M*_*xw*_, *M*_*yw*_, *M*_*zw*_), CEST (*M*_*xs*_, *M*_*ys*_, *M*_*zs*_) and MTC (*M*_*zMTC*_)(1):MTC

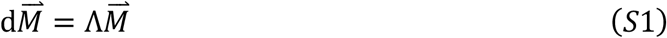

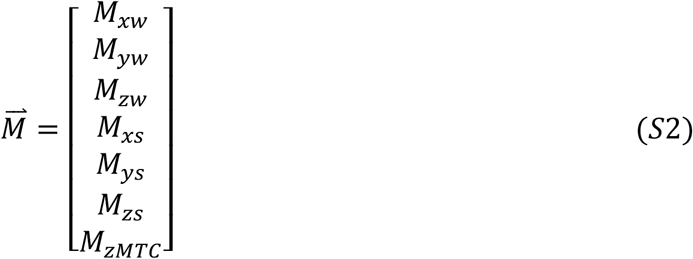

Here, the transverse components of MTC (*M*_*xMTC*_, *M*_*yMTC*_) are set to zero due to the extremely short *T*_*2*_ in the semisolid.

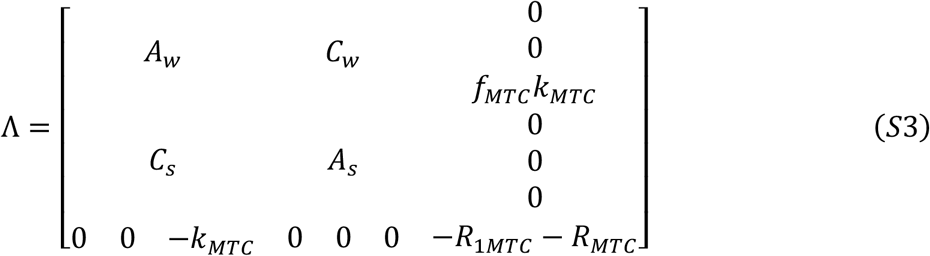

*A*_*w*_ and *A*_*s*_ are two 3×3 matrixes that describe the evolution of the water and solute magnetizations under an RF pulse, respectively, while *C*_*w*_ and *C*_*s*_ are two 3×3 matrixes describing the chemical exchange processes from the water and from the solute pools, respectively. The exchange between solute pool and MTC pool was neglected. *k*_*MTC*_ is the exchange rate from the MTC pool to water, and *f*_*mtc*_ is the fractional concentration of the MTC protons to the water protons. *R*_1*MTC*_ is the longitudinal relaxation rate of the MTC pool. *R*_*MTC*_ is the lineshape of the macromolecular pool together with other saturation transfer components. The MTC lineshape has been reported as Gaussian for Agar and Super-Lorentzian for semi-solid biological tissues (2,3). Considering the frequency range studies in Z-spectra is far smaller than those in MTC studies and the complicated contributions from other exchangeable protons, the MTC lineshape is assumed to be a linear function:

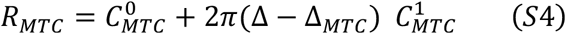

where Δ_*MTC*_ is the offset of the MTC pool resonance center relative to the water signal (Hz). *A*_*w*_ and *A*_*s*_ are given by

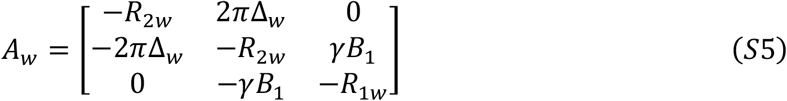

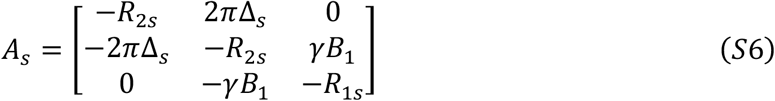

where *R*_2*w*_ and *R*_2*s*_ are the transverse relaxation rates and *R*_1*w*_ and *R*_1*s*_ the longitudinal relaxation rates for the two pools. Δ_*w*_ and Δ_*s*_ are chemical shift offsets (Hz) for the two pools relative to the water proton pool. The exchange matrixes *C*_*w*_ and *C*_*s*_ are given by

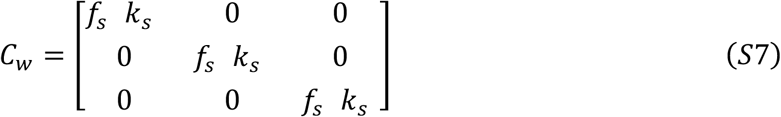

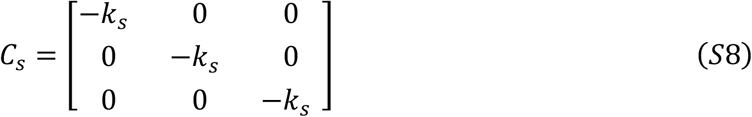

### Exchange rate measurement of the CrCEST

The exchange rates of CrCEST in the cross-linked BSA phantoms were measured by the magnetization recovery of the exchangeable proton peaks after inversion by a single selective Gaussian pulse (width of 2.5 ms) assumed to have negligible effect on the semisolid. After a delay TI, the NMR signal was detected by a 3-9-19 WATER suppression by GrAdient Tailored Excitation (WATERGATE) sequence with a total echo time of 1 ms. Repetition time was 10 s. A list of 15 inversion times from 0 ms to 200 ms were used. The offset was set to 2.0 ppm (with respect to water). The recovery rate was determined by the apparent relaxation rate, 1/*T*_1_ + *k*_*sw*_ ≈*k*_*sw*_, from the fitting of the recovery curve using the following equation

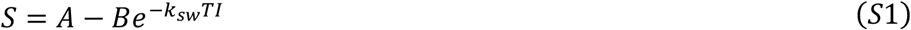

### *In vivo* MRS on GAMT^*-/-*^ mice

The total creatine (Cr + PCr) concentrations in all GAMT-/-mouse brains were obtained from *in vivo* proton MRS experiments performed on voxels of 2.5 × 2.5 × 2.5 mm^3^ each using a stimulated echo acquisition mode (STEAM) sequence (TE = 3 ms, TM = 10 ms, TR = 2.5 s, NA = 64). These MRS spectra for the four GAMT^-/-^ are plotted below and were fitted with the LCModel as detailed previously (4), showing negligible total Cr (1.47±0.68 mM):

**Supporting Information Figure S1:**
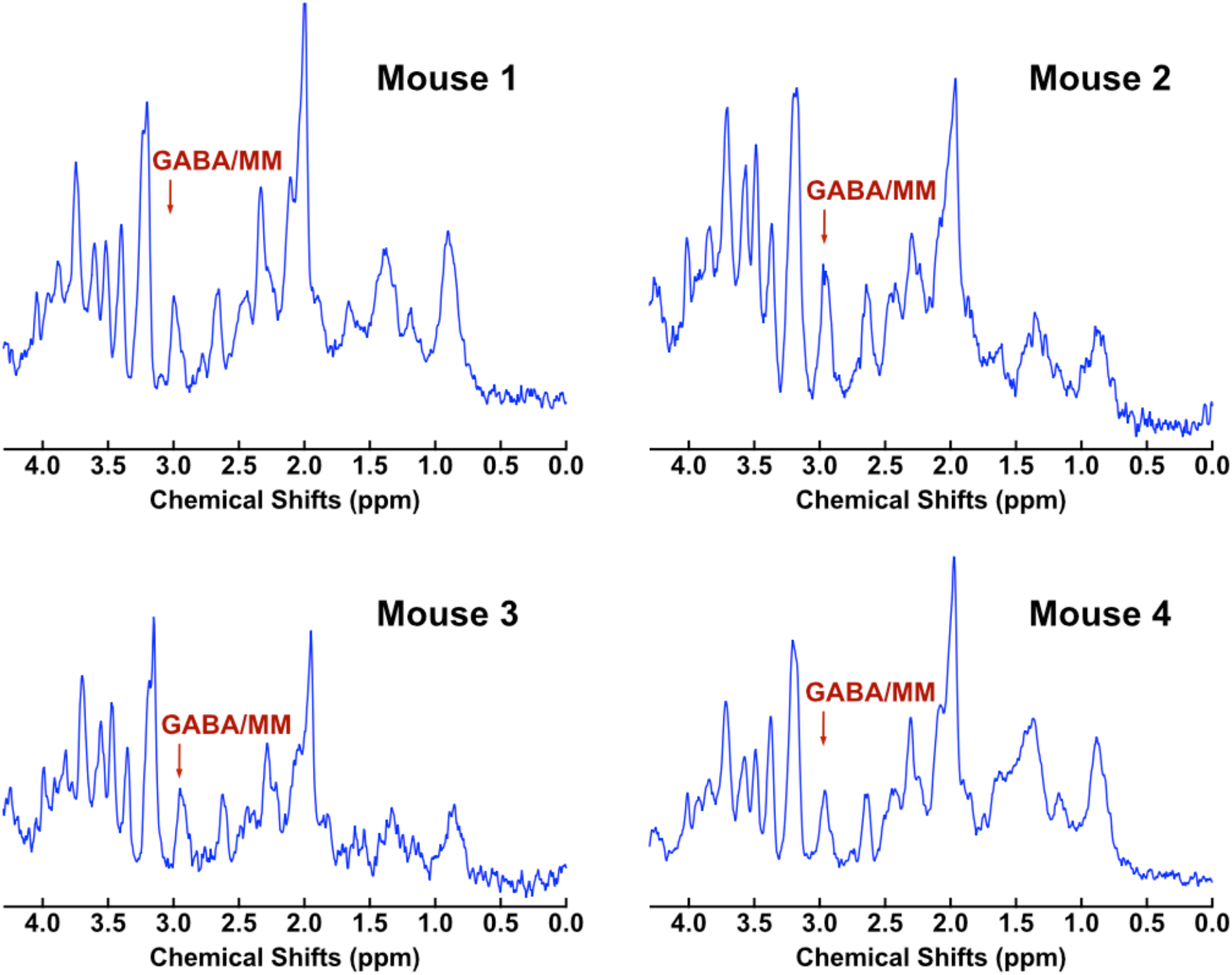
*in vivo* ^1^H NMR spectra measured from regions of the midbrain in all four GAMT^-/-^ mice. The peak at 3 ppm of all GAMT^-/-^ mice shows a residual signal due to gamma aminobutyric acid (GABA) and mobile macromolecules (MM), while LCModel fitting give negligible Cr/PCr remaining.

### Bloch-McConnell (BM) Fitting results

**Table S1:** The *T*_*2app*_,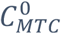 and 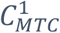 obtained by the MTC background fitting for the CrCrossBSA phantom. *T*_1w_=1.8 s, *T*_1MTC_=1 s, *k*_MTC_= 20 s^-1^ and *f*_*MTC*_ = 10% were fixed during the global BM fitting.

| $B_1$ (μT) | $T_{2app}$ (s) | $C_{MTC}^0$ (s<sup>-1</sup>) | $C_{MTC}^1$ (10<sup>-6</sup>s<sup>-1</sup>/rad) |
| --- | --- | --- | --- |
| 0.4 | 0.0585 | 0.677 | 19.3101 |
| 0.6 | 0.0605 | 0.775 | 11.623 |
| 0.8 | 0.0648 | 1.366 | 15.504 |
| 1.0 | 0.0732 | 2.482 | 42.6371 |
| 1.2 | 0.0862 | 4.143 | 86.290 |
| 1.4 | 0.0994 | 5.461 | 106.096 |
| 1.6 | 0.1089 | 7.073 | 139.772 |

**Table S2:** The *T*_*2app*_,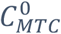 and 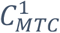 obtained by the MTC and water background fitting for *in vivo* WT mice. *T*_1w_=1.8 s, *T*_1MTC_=1 s, *k*_MTC_= 20 s^-1^ and *f*_*MTC*_ = 10% were fixed during the global BM fitting.

| $B_1$ (μT) | $T_{2app}$ (s) | $C_{MTC}^0$ (s<sup>-1</sup>) | $C_{MTC}^1$ (10<sup>-6</sup>s<sup>-1</sup>/rad)f |
| --- | --- | --- | --- |
| 0.4 | 0.0499 | 0.8106 | 24.920 |
| 0.5 | 0.0173 | 1.118 | 33.726 |
| 0.6 | 0.0285 | 1.803 | 51.731 |
| 0.8 | 0.0298 | 2.898 | 86.045 |
| 1.0 | 0.0303 | 4.119 | 118.428 |
| 1.2 | 0.0310 | 5.173 | 145.000 |
| 1.4 | 0.0309 | 6.173 | 170.193 |
| 1.6 | 0.0382 | 7.592 | 207.347 |

**Table S3:** The *T*_*2app*_,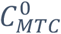 and 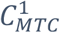 obtained by the MTC background fitting for *in vivo* GAMT^-/-^ mice. *T*_1w_=1.8 s, *T*_1MTC_=1 s, *k*_MTC_= 20 s^-1^ and *f*_*MTC*_ = 10% were fixed during the global BM fitting.

| $B_1$ (μT) | $T_{2app}$ (s) | $C_{MTC}^0$ (s<sup>-1</sup>) | $C_{MTC}^1$ (10<sup>-6</sup>s<sup>-1</sup>/rad) |
| --- | --- | --- | --- |
| 0.4 | 0.0380 | 0.836 | 18.992 |
| 0.5 | 0.0575 | 1.695 | 51.852 |
| 0.6 | 0.0508 | 2.080 | 61.582 |
| 0.8 | 0.0475 | 2.653 | 65.883 |
| 1.0 | 0.0560 | 4.032 | 105.768 |
| 1.2 | 0.0601 | 4.893 | 119.264 |
| 1.4 | 0.0714 | 6.810 | 178.059 |
| 1.6 | 0.0795 | 8.370 | 225.996 |

### Arg and Amide exchange rates in protamine

Protamine has been used previously as model protein for an ArgCEST study (5). Therefore, the exchange rates of guanidinium protons were measured in the protamine solution for comparison with *in vivo* results. Several 50 mg/ml protamine (1.1×10^4^ mM) solutions (Sigma-Aldrich) were prepared in phosphate buffered saline (PBS) and titrated to pH 6.4±0.1, 6.7±0.1, 7.0±0.1, and 7.3±0.1. The protamine solutions were studied in 5 mm NMR glass tubes at 37 °C at 17.6T. The exchange rates of the amide and ArgCEST in protamine were measured with selective inversion recovery method detailed in the main text method section. A list of inversion times (start = 0 ms, end = 60 ms, number of samples = 15) was used for the ArgCEST exchange rate measurement in protamine. The offsets (with respect to water) were set to 3.5 ppm for amideCEST and 2.0 ppm for ArgCEST. 17.6T NMR spectra of protamine for typical pH value are presented in Fig. S2, presenting both amide (3.5 ppm) and Guan (2.0 ppm) exchanging proton groups. Due to the abundant Arg residues in protamine (6), aromatic and other overlapping protons were assumed to be negligible. The Guan peak broadens significantly at pH 7.3, due to the increased exchange rate. Typical inversion recovery curves for the amide and Guan peaks are shown in Fig. S2b, together with the curves fitted using Eq. S1. The pH dependence of the exchange rates for the amide and Guan groups are presented in Fig. S2c. At pH 7.0, *k*_amide_ = 112 ± 5 s^-1^ and *k*_Guan_ = 743 ± 20 s^-1^ were determined. Beyond that, the guan exchange rate is difficult to determine due to the extremely high value, since its saturation/inversion in the selective inversion method will be an obstacle.

**Supporting Information Figure S2:**
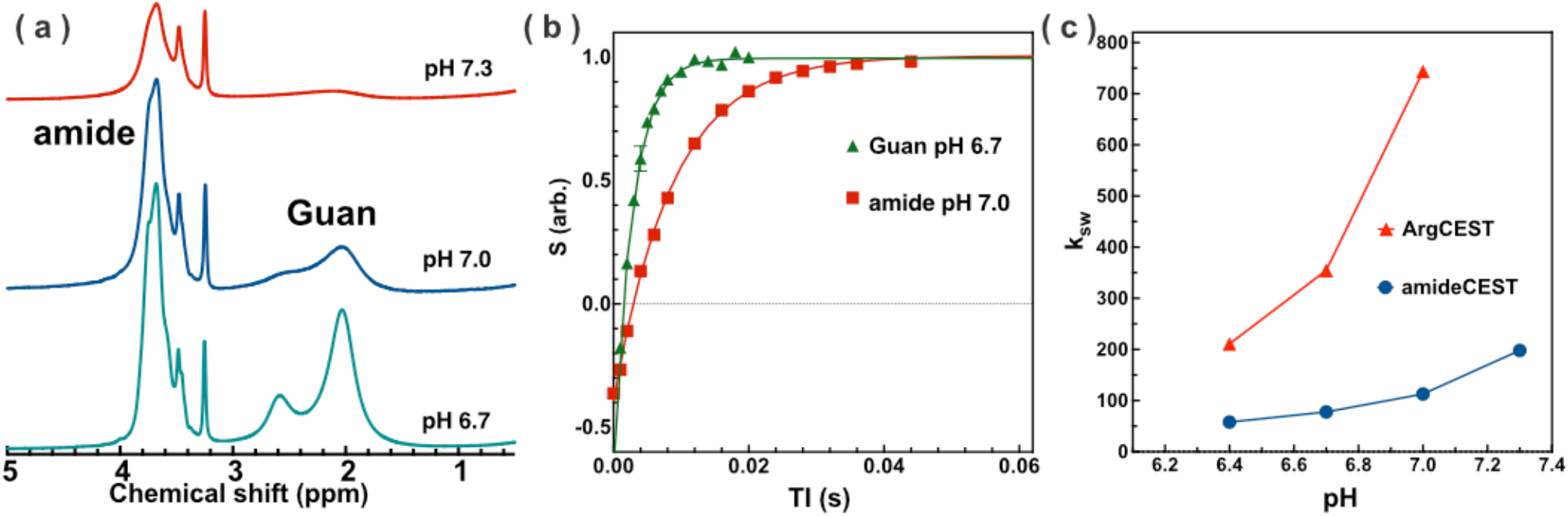
(a) The NMR spectra of protamine solution recorded with a WATERGATE sequence (TE = 1ms) at three pH values (6.7, 7.0 and 7.3). (b) The averaged inversion recovery curves (n=3) of peaks at 2 ppm (Guan) and 3.5 ppm (amide) in protamine solution, respectively. The curves were fitted using a single exponential curve (Eq. 1). (c) pH dependence of the exchange rates for the ArgCEST and amideCEST peaks. All protamine experiments were performed at 37 ºC.

